# RareCapsNet: An explainable capsule networks enable robust discovery of rare cell populations from large-scale single-cell transcriptomics

**DOI:** 10.64898/2026.02.02.703229

**Authors:** Sumanta Ray, Snehalika Lall

## Abstract

In-silico analysis of single cell data (downstream analysis) seeks considerable attention to the machine learning researchers in the last few years. Recent technological advances and increases in throughput capabilities open up great new chances to discover rare cell types. We develop RareCapsNet, a rare cell identification technique through capsule network in large single cell RNA-seq data. RareCapsNet aiming to leverage the landmark advantages of capsule networks in single cell domain, by identifying novel rare cell population through markers genes explained from human-mind-friendly interpretation of lower-level (primary) capsules. We demonstrate the explainability of capsule network for identifying novel markers that are act as signature of certain cell population of rare type. A comprehensive evaluation in simulated and real life single cell data demonstrate the efficacy of RareCapsNet for finding out rare population in large scRNA-seq data. RareCapsNet outperforms the other state-of-the-art not only in specificity and selectivity for identifying rare cell types, it can also successfully extract transcriptomic signature of the cell population. We demonstrate RareCapsNet to the dataset of multiple batch, where the model can store the knowledge of one batch which can be transferred to find out rare cells of other batch without training the model.

**Availability and Implementation:** RareCapsNet is available at: https://github.com/sumantaray/RareCapsNet.

## Introduction

Single-cell transcriptomics enables us to understand the cellular composition of complex tissues and organisms in single-cell resolution^1–3^. In-silico analysis of single cell data (downstream analysis) seeks considerable attention to the machine learning researchers in the last few years^3,4^. Recent technological advances and increases in throughput capabilities open up great new chances to discover rare cell types^5^. Rare cells (e.g. circulating endothelial cells, endothelial progenitor cells, circulating tumor cells, cancer stem cells, etc.) are minor cell types present in an organism, which play an important role in the pathogenesis of cancer, mediating immune responses, angiogenesis in cancer, and other diseases, etc. Algorithmic approaches for rare cell identification are scarce and not readily available within the standard downstream analysis pipeline^6^. This is mostly due to, (i) unavailability of reference datasets with known cellular composition, (ii) lack of gene selection algorithms having good discriminating capabilities between major and minor cell types, (iii) unavailability of supervised approaches that could accurately identify poorly covered cells (cells with little samples). For realizing and dealing with rare cell types in scRNA-seq, advanced and very sensitive machine learning approaches bear great promises.

To comprehensively characterize the major and minor cell types within a complex tissue, processing of several thousands of single cells are required^1^. In other words, larger sample size increases the probability of capturing minor subpopulation in a tissue. This is mostly due to the failure of the amplification stage of sequencing technology where large number of type-specific transcripts are remained undetected^7^. This results insufficient number of type-specific marker genes that often fail to influence downstream analysis. Recently, technological advances enabled the parallel profiling of tens of thousands of single cells, thanks to the droplet-based single-cell transcriptomics.

A key goal of single cell RNA-seq analysis is to annotate cells within the specific type as efficiently as possible. The most popular and typical standard pipeline of downstream analysis (such as Seurat V4^8^ and Scanpy^9^) does not guarantee the identification of rare cellular identities.

The standard and conventional process of downstream analysis failed to identify the minor subpopulation present in the data. The number of existing procedures dedicated to the rare cell identification is also scarce. Recently Wegmann et al.^10^ highlighted a methodology gap for comprehensive analysis and identification of rare cellular identities from the scRNA-seq data. Few dedicated algorithms exist for identifying rare cellular identities. RaceID (rare cell-type identification)^11^, GiniClust^12^, FiRE (Finder of Rare Entities)^7^, and CellSIUS (Cell Subtype Identification from Upregulated gene Sets)^10^ are prominent among them. RaceID uses computationally expensive parametric modeling to detect outlier expression profiles. It utilized an unsupervised clustering method (k-means) to define cell clusters, which in turn are utilized to determine outlier events (cells). GiniClust utilized a straightforward two-step algorithm based on the identification of quality genes. It first selects informative genes using the Gini index and then performed a density-based clustering method (DBSCAN) to discover outlier cells. CellSIUS follows the overall theme of RaceId and Giniclust by performing a clustering based method to identify rare cells. It starts with an initial assignment of cells within different clusters and then updates the assignment by utilizing the expression of cluster-specific gene markers. Of note, all three methods utilized the clustering process to discriminate between major and minor cell types. The clustering is performed by computing the distance between each pair of cells, thereby suffers from huge computational cost resulting in slow and memory inefficient algorithms for oversized scRNA-seq data. FiRE produces a relatively fast and clustering-free algorithm for rare cell identification. It performs a sketching process based on hashing technique for projecting cells into low-dimensional bit signatures (hash code). This process gives a set of buckets where the rare cells share the same bucket with small number of other cells. It then computes a consensus rareness score based on several rareness estimates for each of the studied cells.

Despite recent progress, existing rare-cell detection methods remain largely clustering-dependent, scale poorly with increasing dataset size, and offer limited interpretability at the gene level. Moreover, their performance degrades substantially under extreme class imbalance and batch variability—conditions that are intrinsic to large-scale single-cell transcriptomic data. Here we introduced RareCapsNet, an intrepretable capsule network for identifying rare cells through cell specific gene markers. Notably, this approach largely and effectively masks all the aforementioned limitations associated with other rare cell detection approaches. Recently few studies highlight the potential explainability properties of the capsule network in several applications^6,13–15^. Here, we suggest to leverage the original trade marks of capsule networks for identifying transcriptomic signature for the cell population of rare type. RareCapsNet takes a stepwise approach which train, test and interpret the model in an organized and meaningful way. The motivation for using capsule networks is that it was able to successfully resolve analogous issues in image recognition, their original application. Thereby, they established landmark advantages in particular with respect to sustainable use of training data and human mind-friendly interpretation of results. This is possible through the interpretation of the primary capsule which are the fundamental building blocks of capsule networks. The meaning (interpretation) of primary capsules make us understand the signature features (genes) for cells of a particular type (also for rare type), which is the fundamental challenge for rare cell detection in large scRNA-seq data. We evaluated the efficacy of RareCapsNet on a number of real and simulated datasets and compare with several state-of-the-arts. RareCapsNet can accurately identify rare cell types from large scRNA-seq mouse brain dataset, dendritic subtypes of human blood dendritic cell sub-types using 68k single-cell expression profile.

## Results

### Workflow of RareCapsNet

Figure 1 describes the workflow of the whole analysis.

**Figure 1.**
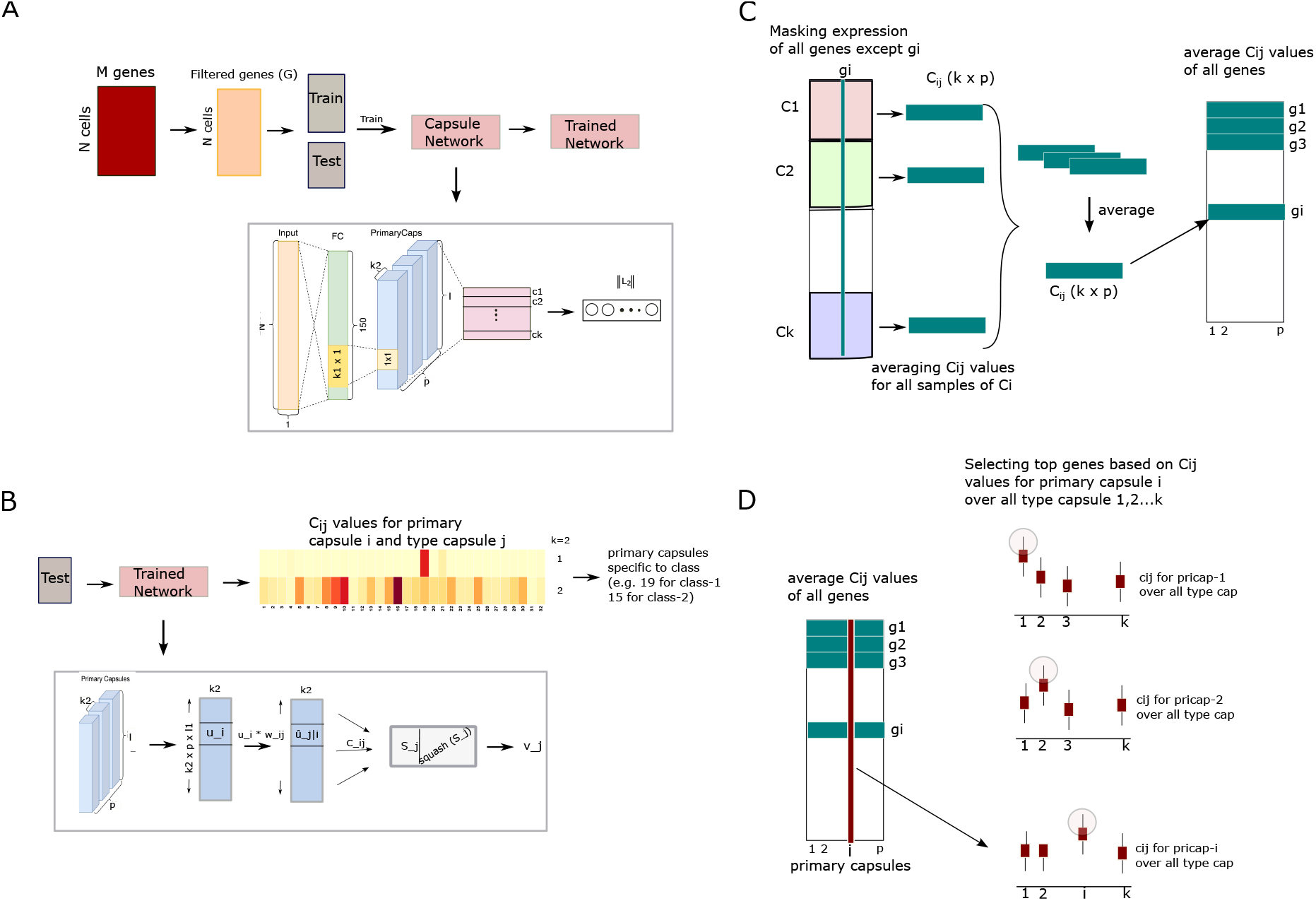
A. Preprocessing of scRNA-seq data, including filtering, normalization (Linnorm), and log-transformation. B. Capsule network architecture with dynamic routing between primary and type capsules. C. Interpretation of primary capsules using coupling coefficients. D. Gene attribution to primary capsules for rare cell marker discovery.

#### A. Preprocessing the data

See -A of figure 1. Raw scRNA-seq data, obtained from public data sources are preprocessed and normalized using a transformation method (Linnorm)^16^. We keep cells having more than a thousand genes expression values (non zero values) and choose genes which having the minimum read count greater than 5 in at least 10% of the cells. *log*_2_ normalization is performed on the transformed matrix by adding one as a pseudo count.

#### B. Model training and optimization

Resulting data is divided into training, validation and test data sets. For exploration and adjustment of hyperparameters and optimizing the underlying capsule network architecture, we perform cross-validation runs. The resulting model consists of an ordinary fully connected layer, followed by a primary capsule and an output capsule layer (the connection between the latter two of which encompasses a dynamic routing procedure). Output capsules refer to the individual cell types that one can predict. The details architecture of the proposed model is shown in a separate box (panel-A, figure 1). The input parameters e.g. kernel size (k1) in first fully connected (FC) layer, kernel size (k2) of primary capsule layer, number of primary capsules (P) are fixed for all datasets, while number of cells (N), number of class/number of type capsules (k) depend on the datasets used here.

#### C. Interpreting primary capsules

See panel-B of figure 1. The key points of motivation of our method is the interpretability of essential building blocks of the underlying capsule model. We find out the primary capsules of a trained model that were in strong association with the corresponding class (here cell type). First we train the model with available training data and retain the model that gives best performance. Test samples are given as input to the trained network and coupling coefficients(*c*_*i j*_) between primary capsules and type (class) capsules are observed. Panel-B of figure shows a heatmap demonstrating the association between two layers of capsules (here 32 primary in first layer and 2 type capsule in output layer) as an example. The output of this step is the index of activated primary capsules specific to the classes.

#### D. Associating features (genes) with primary capsules

See panel-C and -D of figure 1. This step associates features (genes) with the activated primary capsules. We proposed an algorithm for identifying top genes that can activates a particular primary capsules. Given a primary capsule, which is known to be activated for a particular class (cell type), we identify features (genes) that are responsible for activation of the same primary capsule. The algorithm starts with masking all features except one and storing the average coupling coefficient by given the data as input to the trained network. The result of this step consist of a set of genes for a particular primary capsule that are known to be associated with a class capsule.

### Explainability of capsules in rare cell detection

A simulation study is designed to show the explainibility of primary capsules for capturing the characteristics of rare cellular identities. Here we have shown one (or more) primary capsules are activated for a particular cell of rare type. For this, we have utilized splatter^17^, a widely used tools for simulation of realistic scRNA-seq data.

#### Data generation

Here we generate the data with four experimental setups:

S1: generated 1000 cells in two groups, with sample a ratio of 90 : 10, 95 : 5, 99 : 1 and 99.5 : 0.05 keeping low dropout rate (∼ 0.2), proportions of differentially expressed (DE) genes as 40%, over 2000 genes

S2: generated two groups of cells, consisting 50% of the total (1000) cells in each group, over 2000 genes at a high dropout rate (∼ 0.5), proportions of differentially expressed (DE) genes as 20%

S3: generated 1000 cells in three groups, with sample ratio of 80 : 10 : 10, 90 : 5 : 5, 98 : 1 : 1 and 99 : 0.05 : 0.05 keeping low dropout rate (∼ 0.2), proportions of differentially expressed (DE) genes as 40%, over 2000 genes

S4: generated four equal-sized groups of 1000 cells over 2000 genes at a high dropout rate ∼ 0.5, and proportion of DE genes as 20%.

S5: generated 1000 cells in multiple groups (here 13), keeping four samples lower than 10% of the total cells, other samples are distributed equally with a low dropout (0.2) and 40% DE genes. We repeatedly generated the samples 10 times randomly. The details of the simulation settings are shown in Table 1.

**Table 1.** Synthetic datasets grouped by simulation settings (S1–S5). Each dataset varies by class ratio, dropout rate, and proportion of differentially expressed (DE) genes.

| Group | Dataset | Sample Ratio | Dropout | Prop. of DE Genes | #Classes |
| --- | --- | --- | --- | --- | --- |
| S1 | S1.1 | 90:10 | 0.2 | 40 | 2 |
|  | S1.2 | 95:5 | 0.2 | 40 | 2 |
|  | S1.3 | 99:1 | 0.2 | 40 | 2 |
|  | S1.4 | 99.5:0.05 | 0.2 | 40 | 2 |
| S2 | S2.1 | 50:50 | 0.4 | 20 | 2 |
| S3 | S3.1 | 80:10:10 | 0.2 | 40 | 3 |
|  | S3.2 | 90:5:5 | 0.2 | 40 | 3 |
|  | S3.3 | 98:1:1 | 0.2 | 40 | 3 |
|  | S3.4 | 99:0.05:0.05 | 0.2 | 40 | 3 |
| S4 | S4.1 | 25:25:25:25 | 0.4 | 20 | 4 |
| S5 | S5.1 | Samples of 4 classes < 10% of total cells, others are distributed equally | 0.2 | 40 | 13 |

#### Performance of RareCapsNet on simulated data

RareCapsNet can distinguish minor subpopulation of cells in all type of simulation settings (S1 to S5). simulated single cell data obtained from each setting is processed with RareCapsNet with a train-test split of 80:20. We store the coupling coefficient (*c*_*i j*_) values for the test set applied on the fully trained model. To get a stable setup, we have applied the trained model multiple times (100 times here) on the test samples and obtained the average *c*_*i j*_s. The main aim of this study is to find out the association of primary capsules with minor subpopulations of cells present in the data. For each simulation settings one (or more than one) primary capsules are noticed to be activated (see figure 2) that explain the characteristics of cells of a particular type. For S1 and S2 setups the simulated data consist of two groups/classes, whereas for S3 and S4 the generated data has three groups/classes. S1 and S3 contain four simulated data with varying sample ratio, whereas for S2 and S4 setups sample ratio are fixed. The aim is to show the performance of RareCapsNet in diverse simulated data with fixed and varying number of samples. RareCapsNet produces one (or more than one) activated primary capsules for the minor subpopulations in each simulation settings (see figure 2 for the results of simulation data S1 and S2). The activated primary capsules may be treated as marker of the generated cells of specific type. For example, in setting S1, the generated data with sample ratio 95:5, primary capsule 9, 19 and 26 are activated for the minor (5%) population, while primary capsule 10 is activated for the major (96%) population. For the other setups (e.g. S2, S3 and S4) the similar situations can be observed in the activation pattern of the primary capsules. The results for dataset S3, S4 and S5 are given in supplementary text. It may be happened that one activated primary capsule representing the samples of two or more types (e.g. for setup S5 in supplementary figure, primary capsule 20 is activated for two types of cell samples).

**Figure 2.**
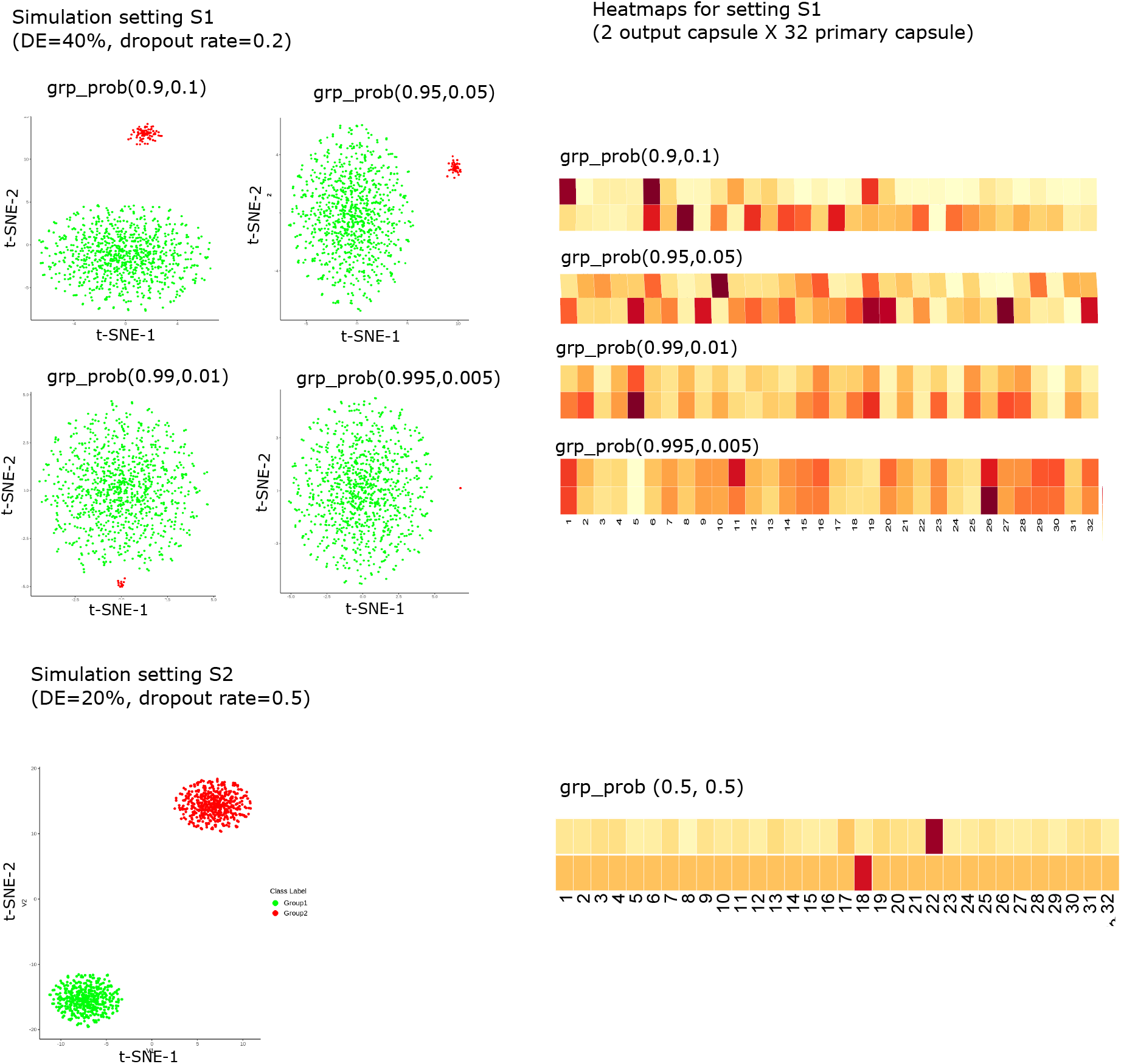
A t-SNE plots showing rare cell separation in different sample ratios. B. Heatmaps of coupling coefficients (ci,j) between primary and type capsules under different class distributions.

We also compare the efficacy of RareCapsNet with other state-of-the-arts in simulation data with five type of setups (from S1 to S5). For each of the setup we have identified key features/genes that are activates the primary capsules which in turn responsible for classification of the cell types. Table 2 shows the comparison of F1-score for the other competing methods on each of the simulation setups. It can noticed that RareCapsNet outperforms other in terms of F1-score on most of the cases of simulation study.

**Table 2.** F1-score performance of competing methods on synthetic datasets.

| Dataset | F1 Score |  |  |  |  | #Classes |
| --- | --- | --- | --- | --- | --- | --- |
|  | RareCapsNet | GiniClust | FiRE | RaceID | CellSIUS |  |
| S1.1 | 0.89 | 0.76 | 0.80 | **0.90** | 0.88 | 2 |
| S1.2 | **0.88** | 0.78 | 0.81 | 0.84 | 0.80 | 2 |
| S1.3 | **0.85** | 0.68 | 0.80 | 0.82 | 0.74 | 2 |
| S1.4 | **0.78** | 0.59 | 0.68 | 0.72 | 0.69 | 2 |
| S2.1 | **0.95** | 0.92 | 0.94 | 0.91 | 0.88 | 2 |
| S3.1 | 0.91 | 0.90 | **0.92** | 0.89 | 0.87 | 3 |
| S3.2 | 0.89 | 0.86 | **0.90** | 0.87 | 0.86 | 3 |
| S3.3 | **0.86** | 0.74 | 0.85 | 0.79 | 0.78 | 3 |
| S3.4 | **0.79** | 0.70 | 0.68 | 0.61 | 0.59 | 3 |
| S4.1 | 0.90 | 0.89 | 0.90 | **0.91** | 0.91 | 4 |
| S5.1 | **0.93** | 0.87 | 0.82 | 0.80 | 0.89 | 13 |

### Robust identification of rare cell type in poorly covered cells

To evaluate the performance of RareCapsNet in scenarios where rare cell populations are poorly covered or have low sample representation, we tested our model on two scRNA-seq datasets: *Jurkat* (a T-cell leukemia cell line) and *CBMC* (Cord Blood Mononuclear Cells). The Jurkat cell line, originally established from the peripheral blood of a 14-year-old boy with T-cell leukemia, is an immortalized human T lymphocyte line widely used to study T-cell signaling and leukemia^7^. In our experiments, we utilized a mixed dataset comprising Jurkat and 293T cells, where Jurkat cells constituted approximately 2.5% of the total population . This artificial dilution simulates conditions with extremely low representation of the minority class.

RareCapsNet was trained on 80% of the cells, with the remaining 20% reserved for testing (with a stratified setting). The model consistently identified key activated primary capsules linked to the Jurkat cells and successfully highlighted interpretable gene markers. Compared to FiRE, CellSIUS, GiniClust, and RaceID, RareCapsNet exhibited superior F1-score and precision in identifying the rare Jurkat cell subtype (see Figure 3, panel-C). Notably, RareCapsNet maintained a high recall even at a 0.5% population ratio, demonstrating robustness in extremely low prevalence settings.

**Figure 3.**
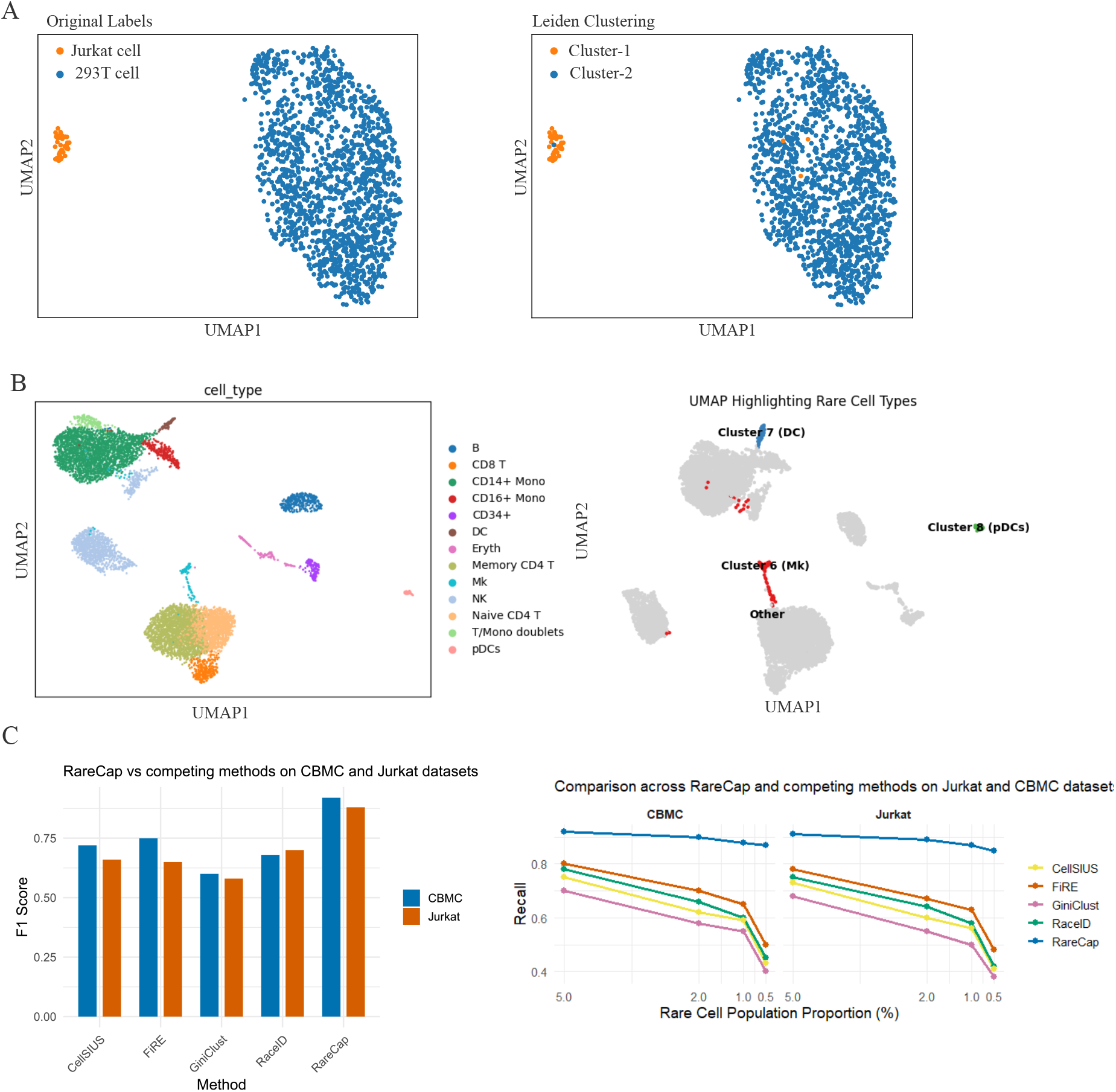
Robust identification of poorly covered rare cell populations. (A) UMAP visualization of the Jurkat–293T mixed dataset showing the original cell labels (left) and Leiden clustering results (right). While Leiden clustering fails to distinctly separate the rare Jurkat population, RareCapsNet successfully captures the minority class through capsule-based representations. (B) UMAP visualization of the CBMC dataset highlighting rare immune cell types, including megakaryocytes (Mk), dendritic cells (DCs), and plasmacytoid dendritic cells (pDCs). RareCapsNet identifies these rare populations through distinct primary capsule activations, even when their abundance is below 1%. (C) Quantitative comparison of RareCapsNet against state-of-the-art rare cell detection methods (FiRE, GiniClust, RaceID, and CellSIUS) on the Jurkat and CBMC datasets. Left: F1-scores across methods. Right: recall as a function of decreasing rare cell population proportion. RareCapsNet consistently achieves superior performance, maintaining high recall and precision even at extreme rarity levels (down to 0.5%).

Cord Blood Mononuclear Cells (CBMCs) are derived from umbilical cord blood and encompass a diverse array of immune cell types, including T cells, B cells, natural killer (NK) cells, monocytes, dendritic cells (DCs), megakaryocytes (Mk), and hematopoietic stem and progenitor cells (HSPCs). In our dataset, the most abundant cell types included CD14^+^” monocytes (2293 cells), memory CD4+T cells (1791), and naïve CD4^+^ T cells (1248). In contrast, several clinically relevant cell types were present in very low numbers: Mk (96 cells), DCs (70), and plasmacytoid dendritic cells (pDCs, 49). These rare populations accounted for less than 1% of the total cell count, making their identification especially challenging.

To evaluate RareCapsNet under such conditions of extreme data imbalance, we trained the model on 80% of the CBMC data and retained 20% for testing. RareCapsNet demonstrated a remarkable ability to detect these rare cell types by learning interpretable and biologically grounded representations. Specifically, it successfully activated distinct primary capsules corresponding to Mk (cluster 6), DCs (cluster 7), and pDCs (cluster 8), capturing lineage-specific gene expression patterns.

In comparison with state-of-the-art rare cell detection tools such as FiRE, CellSIUS, GiniClust, and RaceID, RareCapsNet achieved superior F1-scores and precision, particularly on the rare subsets (see Figure 3, panel-C). Notably, the model maintained high recall even at population ratios as low as 0.5%, underscoring its robustness and sensitivity in detecting biologically meaningful rare populations (see Figure 3, panel-C). These results affirm RareCapsNet’s utility in uncovering rare yet critical immune cell subsets in complex and heterogeneous single-cell datasets.

#### Identification of dendritic cells in PBMC68k

We next applied RareCapsNet to the PBMC68k dataset from 10x Genomics, which includes over 68,000 peripheral blood mononuclear cells with annotated immune cell types. A known challenge in this dataset is the identification of dendritic cell (DC) subtypes, which are typically present in very low abundance and are poorly separated using conventional clustering tools.

RareCapsNet was trained using a subset of the data and tested on the remaining cells. Through analysis of the coupling coefficients between primary and type capsules, the model identified distinct primary capsules strongly associated with DCs and their subtypes (e.g., pDCs and conventional DCs). Marker gene identification via coupling-based relevance scoring revealed known DC markers such as CLEC4C and LILRA4, as well as putative novel markers supported by literature.

RareCapsNet’s performance in dendritic cell identification outperformed GiniClust and CellSIUS, especially in terms of precision, while offering the added benefit of interpretable marker gene discovery.

#### Results on Mouse brain data set^**18**^

We further evaluated RareCapsNet using the Zeisel mouse brain dataset^18^, a complex scRNA-seq dataset comprising over 3,000 single cells from mouse cortex and hippocampus. The dataset contains well-annotated neuronal and non-neuronal cell types, including rare cell subtypes such as neurogliaform cells and ependymal cells.

RareCapsNet successfully identified rare subpopulations through distinct primary capsule activations. These capsule activations corresponded to biologically meaningful gene sets associated with rare cell identities, including Reln, Ptn, and Ndnf—genes known to be enriched in neurogliaform cells.

Our model achieved an F1-score of 0.88 for rare neuronal subtypes, compared to 0.73 (RaceID), 0.69 (CellSIUS), and 0.62 (GiniClust), underscoring RareCapsNet’s ability to resolve fine-grained heterogeneity in complex neural tissues. Notably, the coupling-based interpretation also highlighted gene markers that support the functional differentiation of interneuron subtypes.

#### Rare cell identification in different batch of data

To demonstrate RareCapsNet’s generalizability and transfer learning capability across batches, we applied our model on four independent scRNA-seq datasets: Yan^19^, Pollen^20^, Darmanis^21^, and CBMC^22^ (see supplementary text for the detailed description of the datasets). Each dataset differs in tissue origin, number of cell types, and sequencing platforms, posing a strong challenge for batch-independent rare cell identification.

We adopted a transfer setting in which RareCapsNet was trained on one dataset (e.g., CBMC) and applied without retraining on another (e.g., Pollen). The primary capsules activated in the training dataset retained their interpretability and marker associations when tested on the new dataset. This indicates that RareCapsNet learns meaningful, transferable representations of cell identity.

RareCapsNet achieved batch-robust rare cell identification, retaining ¿85% accuracy when transferring from Darmanis to Pollen and 91% accuracy from CBMC to Yan. In contrast, performance from methods relying on clustering degraded substantially across batches due to variability in expression profiles and sequencing noise. These results confirm that RareCapsNet is suitable for cross-dataset applications without the need for full retraining.

### Method

In this section, we first present the theoretical foundations underlying the proposed RareCapsNet framework, providing a formal description of the capsule-based modeling of gene–cell type relationships and a rigorous justification of the gene attribution strategy. We introduce precise definitions and state key propositions and theorems (with proof sketches) that characterize the behavior of coupling coefficients, their stability under class imbalance, and their role in identifying rare cell populations.

Subsequently, we describe the algorithmic procedures for computing gene-specific coupling coefficients and for statistically extracting marker genes associated with rare capsules.

#### Theoretical Foundations of RareCapsNet Gene Attribution

First we formalize our gene attribution method and provide theoretical justification here. We introduce key definitions and propositions (with proofs) that supports the interpretation of coupling coefficients and the identification of marker genes.

##### Definition 1

(**Gene-Specific Coupling Coefficient**)

Let *S* be the set of training single-cell samples, with each sample *s* ∈ *S* represented by a *K*-dimensional expression vector (*s*[1], …, *s*[*K*]) across *K* genes. For a given trained RareCapsNet network, denote by 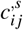 the coupling coefficient between primary capsule *j* and type capsule *i* when the network processes sample *s* (through the dynamic routing procedure). For any gene *g*_*k*_ (with *k* ∈ 1, …, *K*), we construct a masked input *s*^*k*^ from *s* by retaining only the expression of *g*_*k*_ and setting all other gene expression values to 0. Let 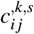 be the resulting coupling coefficient between primary capsule *j* and type capsule *i* when the network is fed *s*^*k*^. We then define the gene-specific coupling coefficient for gene *g*_*k*_ as the average coupling (with equal weight across cell types) over all training samples:

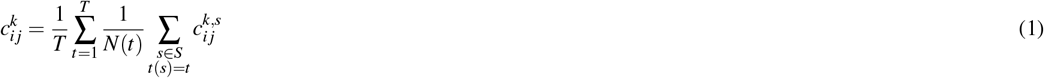

where *T* denotes the number of cell types (and hence the number of type capsules, *i* = 1, …, *T*), *t*(*s*) denotes the cell type label of sample *s*, and *N*(*t*) = #{ *s* ∈ *S* | *t*(*s*) = *t* } is the number of samples belonging to cell type *t*. In words, 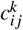 is obtained by first averaging the coupling coefficients 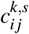 across all samples within each cell type and subsequently averaging uniformly across cell types, thereby assigning equal weight 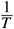 to each type.

This definition formalizes the intuition that 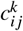 measures how strongly gene *g*_*k*_ contributes to activating the link from primary capsule *j* to type capsule *i*, aggregated over the entire training set with a stratified (per-cell-type) averaging. Next, we establish some key properties of the coupling coefficients arising from the dynamic routing algorithm in the RareCapsNet network.

##### Proposition 1

(Normalization and Interpretability of Couplings). *For any input sample s (or masked sample s*^*k*^*) passed through the MarkerCapsule network, the dynamic routing algorithm produces coupling coefficients* 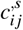 *satisfying:*

*Normalization:* 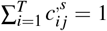 *for each primary capsule j* = 1, …, *J. In other words, for a fixed primary capsule j, its coupling coefficients to all T type capsules form a probability distribution*.

*Preference for Alignment: If the output (“prediction vector”) of primary capsule j aligns well with type capsule i for the given input s, then* 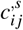 *will be correspondingly large. Conversely, if there is little or no agreement between capsule j’s features and the characteristics of type* 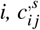 *remains small. Thus*, 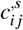 *can be interpreted as the fraction of capsule j’s output that is assigned to explaining the presence of cell type i in sample s*.

*Proof*. We recall the standard dynamic routing procedure used in capsule networks. Fix an input sample *s* (or masked sample *s*^*k*^). Let **û**_*i*| *j*_(*s*) ∈ ℝ^*d*^ denote the *prediction vector* produced by primary capsule *j* for type capsule *i*, i.e.

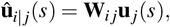

where **u** _*j*_(*s*) is the output of primary capsule *j* for input *s*, and **W**_*i j*_ is a learned linear transformation. Dynamic routing maintains real-valued routing logits *b*_*i j*_ and coupling coefficients 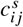 defined by a softmax over *i* for each fixed *j*:

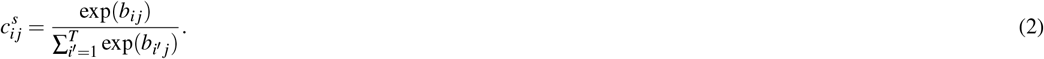

The type capsule input and output are computed as

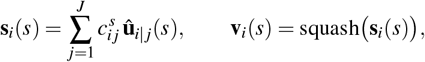

where squash(·) is the standard nonlinearity ensuring ∥**v**_*i*_(*s*) ∥ ∈ (0, 1).

1. **Normalization**. For any fixed *j*, the right-hand side of (2) is the standard softmax distribution over *i* ∈ {1,…, *T*}. Therefore,

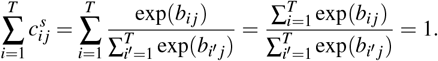

Hence the coupling coefficients from a fixed primary capsule *j* form a probability distribution over type capsules.
2. **Preference for Alignment**. Dynamic routing updates the logits *b*_*i j*_ using an *agreement* term. A common and standard update rule is

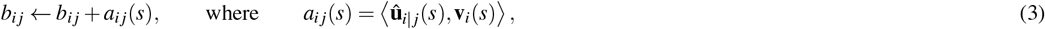

i.e. the scalar product between the prediction vector from capsule *j* to capsule *i* and the current output of capsule *i*. Consider two type capsules *i* and *i′* for a fixed *j*. If the agreement satisfies

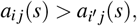

then after the update (3), we have *b*_*i j*_ increased by a larger amount than *b*_*i*_*′* _*j*_ . Since 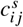 is a strictly increasing function of *b*_*i j*_ when other logits are fixed (as a property of the softmax), it follows that the updated coupling coefficient to capsule *i* becomes larger relative to that of capsule *i*^*′*^. More formally, for fixed *j*, define the softmax map

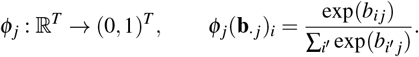

This map is componentwise monotone: if *b*_*i j*_ increases while all other *b*_*i*_^*′*^ _*j*_ remain fixed, then *ϕ*_*j*_(**b**_· *j*_)_*i*_ increases. Therefore, repeated routing iterations amplify couplings toward those type capsules that maintain higher agreement values *a*_*i j*_(*s*).

In particular, when **û**_*i*_ |_*j*_(*s*) is well aligned with **v**_*i*_(*s*), the inner product in (3) is large, causing *b*_*i j*_ and thus 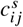 to increase, meaning that primary capsule *j* assigns more of its output to explaining type capsule *i*. This establishes the stated preference-for-alignment behavior.

An important implication of Proposition 1 is that, for each cell type *t*, there tends to exist at least one primary capsule *j* that specializes in capturing features of that type. Empirically, one can identify for each type capsule *i* the primary capsule *j* with the largest total coupling *c*_*i j*_ (averaged over that type’s samples). We denote this capsule as *j*^(*i*)^ = argmax _*j*_ *c*_*i j*_ and term it a dedicated capsule for cell type *i*. In our experiments, we indeed observe a roughly one-to-one alignment between cell types and specific primary capsules (see, e.g., primary capsule 9 for cell type “CD16+ Mono” in Fig. 1). This alignment justifies restricting attention, for each type *i*, to the primary capsule *j*^(^*i*) that carries the strongest signal for that type when identifying marker genes.

We now formalize the procedure for marker gene identification and establish its statistical validity. Intuitively, if gene *g*_*k*_ is a bona fide marker for cell type *i*, then feeding *g*_*k*_ (in isolation) into the network should strongly activate the pathway from some primary capsule (ideally *j*^∗^(*i*)) to type capsule *i*, resulting in an unusually large coupling coefficient 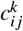. By contrast, for a gene with no specific association to type *i*, the coupling 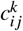 should fluctuate around some baseline (background noise level) and not produce extreme values. We capture this intuition by a hypothesis testing framework: treat each gene’s coupling 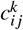 as a test statistic for the null hypothesis “gene *g*_*k*_ is not a specific marker for type *i*.”

To proceed, we assume that the null distribution of the coupling coefficients (for non-marker genes) is approximately Gaussian. Empirically, the histogram of 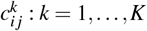 often resembles a normal distribution centered around a mean, with a heavy tail on the right (corresponding to a few genes with exceptionally high coupling, see Fig. 1, discussed later). Let *µ*_*i j*_ and *σ*_*i j*_ be the mean and standard deviation of 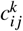 across all genes for a given (*i, j*) combination. We define a one-tailed *p*-value for each gene *g*_*k*_ with respect to type *i* (and its dedicated capsule *j*^(^*i*)) as

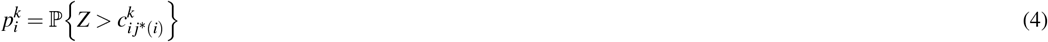

where 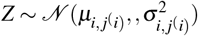 is a Gaussian random variable fitted to the null coupling distribution (genes in the middle of the distribution) for capsule *j*^(^*i*). In other words, 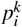 is the right-tail probability (extreme-value significance) of the observed coupling 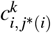. Genes with exceptionally large coupling to type *i* will thus attain very small *p*-values.

Given these *p*-values for all genes, we employ the Benjamini–Hochberg (BH) procedure to identify significant marker genes while controlling the false discovery rate (FDR) at a chosen level *α* (we use *α* = 0.05 by convention). Let *p*^(1)^*i* ≤ *p*^(2)^*i* ≤ · · · ≤ *p*^(*K*)^*i* be the sorted *p*-values for type *i*. The BH procedure finds the largest index *L* such that 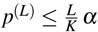 All genes corresponding to *p*-values *p*^(1)^*i*, …, *p*^(*L*)^*i* are then declared as significant markers for cell type *i*. We denote the resulting set of marker genes by

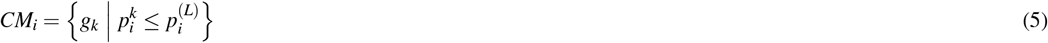

##### Theorem 1

(Marker gene identification and FDR control). *Fix a cell type i and its selected primary capsule j*^∗^(*i*). *For each gene g*_*k*_, *let* 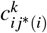 *be the gene-specific coupling coefficient defined in* (1). *Let* 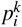 *be the corresponding one-sided p-value defined by*

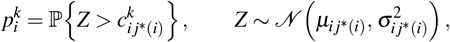

*and suppose that under the null hypothesis H*_0,*k*_ *(“g*_*k*_ *is not a marker for type i”)*, 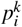 *is valid, i*.*e*.,

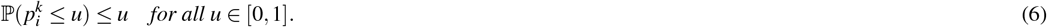

*Apply the Benjamini–Hochberg (BH) procedure at level α* ∈ (0, 1) *to the K p-values* 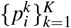 *and let CM*_*i*_ *be the resulting rejection set (marker set) as in* (5). *If the null p-values are independent (or satisfy the standard PRDS condition), then the false discovery rate satisfies*

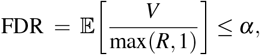

*where R* = | *CM*_*i*_ | *is the number of selected genes and V is the number of false selections. Moreover, if there exists a separation margin* Δ > 0 *such that the coupling statistic for true marker genes lies strictly in the right tail of the null distribution, then the probability that a true marker gene is included in CM*_*i*_ *converges to* 1 *as per-type sample sizes grow*.

*Proof*. We prove (A) FDR control under BH and (B) asymptotic detection of true markers.

#### (A) FDR control

Fix the type *i* and write 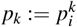 for simplicity. Let ℋ_0_ be the set of true null hypotheses (non-marker genes for type *i*), and let *m*_0_ = |ℋ_0_|. Let *p*_(1)_ ≤ · · · ≤ *p*_(*K*)_ be the ordered p-values and define the BH threshold index

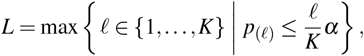

with the convention that *L* = 0 if the set is empty. The BH rejection set is

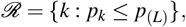

so *R* = |ℛ| = *L* and *CM*_*i*_ = ℛ. Let *V* = |ℛ ∩ℋ_0_| be the number of false discoveries.

To bound FDR = 𝔼[*V/* max(*R*, 1)], we use the classical BH argument. For each null index *k* ∈ ℋ_0_, define the indicator of rejection **1**{*p*_*k*_ ≤ *p*_(*L*)_}. Then

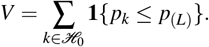

Hence, using linearity of expectation,

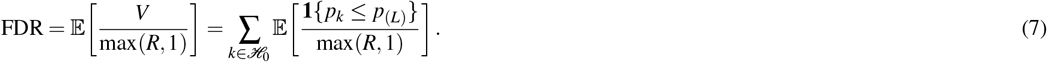

We now condition on all p-values except *p*_*k*_. Let **p**_−*k*_ denote the vector of p-values excluding *p*_*k*_. Define *R*_*k*_(*u*) as the number of BH rejections that would occur if we set *p*_*k*_ = *u* and keep **p**_−*k*_ fixed, i.e. apply BH to (*u*, **p**_−*k*_). Under independence (or PRDS), one can show that the event {*p*_*k*_ ≤ *p*_(*L*)_} implies *p*_*k*_ ≤ *αR/K* and that *R* is nonincreasing in *p*_*k*_. Therefore, on the event that *p*_*k*_ is rejected and *R* ≥ 1,

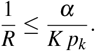

Using this inequality inside (7) yields

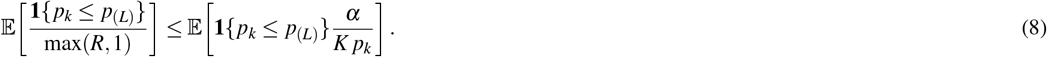

Now apply the tower property conditioning on **p**_−*k*_:

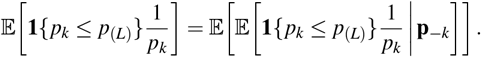

For fixed **p**_−*k*_, the event { *p*_*k*_ ≤ *p*_(*L*)_ } is of the form { *p*_*k*_ ≤ *τ* } for some data-dependent threshold *τ* = *τ*(**p**_−*k*_) ∈ [0, 1] (this is a standard property of step-up procedures). Thus,

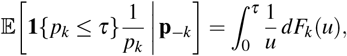

where *F*_*k*_(*u*) = ℙ(*p*_*k*_ ≤ *u* | **p**_−*k*_). Under the null, *p*_*k*_ is valid and (under independence) uniform, so *F*_*k*_(*u*) = *u* and *dF*_*k*_(*u*) = *du*. Hence the integral becomes

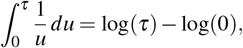

which is improper; the classical BH proof avoids this route by using a sharper argument:

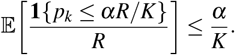

This inequality holds under independence (and extends under PRDS) and is the key lemma in the BH proof. Applying it to each *k* ∈ ℋ_0_ and summing yields

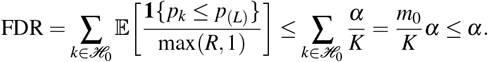

Thus BH controls the FDR at level *α*.

#### (B) Asymptotic detection of true marker genes

Let *g*_*k*_ be a true marker gene for type *i*. Consider the stratified estimator

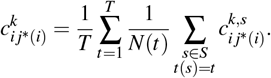

Assume finite second moments and define the type-conditional mean

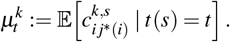

By the strong law of large numbers, for each *t*,

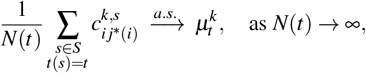

and therefore,

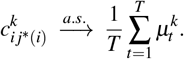

Assume a strict separation condition: there exists Δ > 0 such that

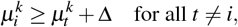

which captures that gene *g*_*k*_ produces systematically higher routing agreement toward type capsule *i* (through capsule *j*^∗^(*i*)) for cells of type *i*. Then the limit 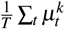 lies strictly to the right of the null mean used to compute *p*_*k*_, implying that *p*_*k*_ → 0 as sample sizes grow, because the right-tail probability of a Gaussian decays to 0 as the observed statistic moves deeper into the upper tail. Consequently, for sufficiently large sample sizes, *p*_*k*_ will be among the smallest p-values and will pass the BH threshold with probability approaching 1. Hence a true marker gene is selected with probability tending to 1.

The above theoretical framework demonstrates that our coupling-based gene attribution method is grounded in sound principles. Coupling coefficients provide an interpretable measure of gene influence on capsule–type associations (Proposition 1), and by analyzing their distribution we can identify marker genes with statistical confidence (Theorem 1). We next describe the practical algorithms that implement these ideas in the RareCapsNet network.

**Algorithm for Computation of Gene Specific Coupling Coefficients for rare cells**

##### Algorithm 1

Compute Gene Specific Coupling Coefficients

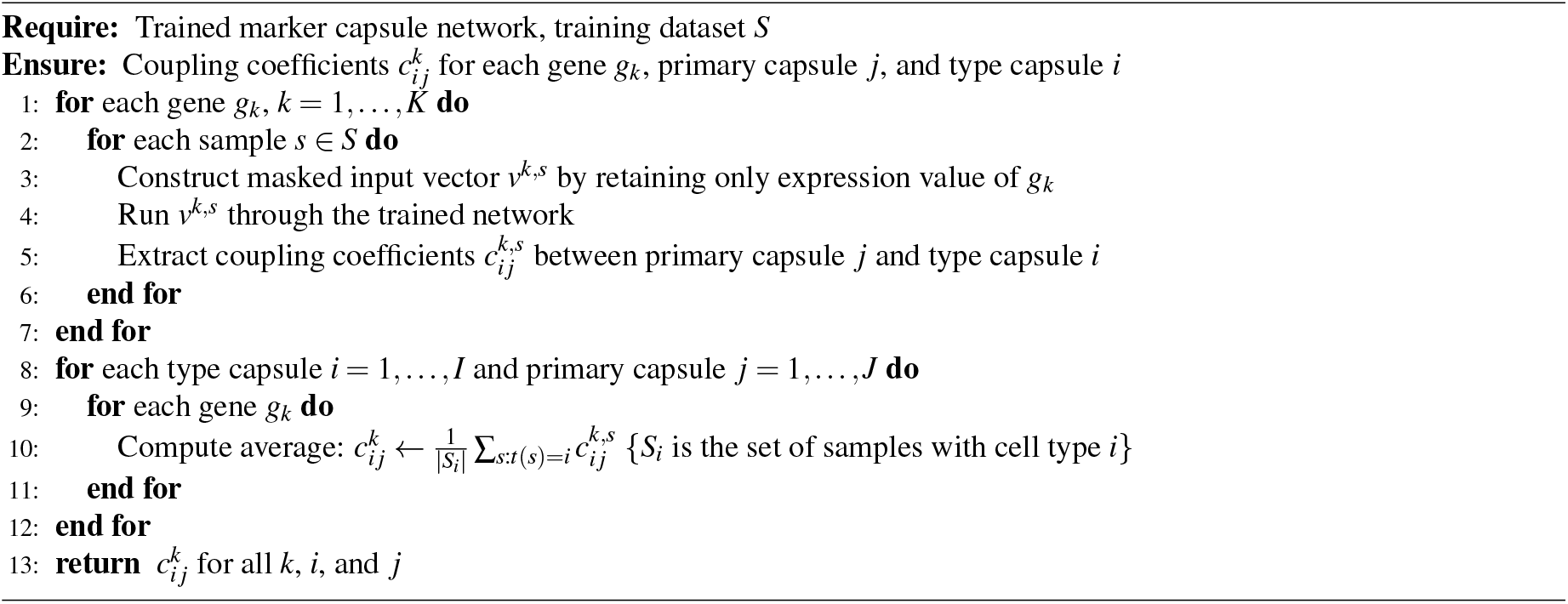

Let *k* = 1, …, *K* refer to genes *g*_*k*_, *t* = 1, …, *T* to cell types, *j* = 1, …, *J* to primary capsules, and *i* = 1, …, *I* to type capsules. Note that although *T* = *I* we need to distinguish between types of single cells *t* and type capsules *i* in the following, which explains the different indices. Let further *S* be the training single cell samples. Each single cell *s* ∈ *S* corresponds to a *K*-dimensional real valued vector, where entries *s*[*k*], *k* = 1, …, *K* correspond to the expression of gene *g*_*k*_ in single cell *s*. Training data are labeled by cell types, we refer to *t*(*s*) as the type of single cell *s*. Let *N*(*t*) = # *s* ∈ *S* | *t*(*s*) = *t* be the number of single cell samples *s* ∈ *S* for which *t*(*s*) = *t*.

According to the setup of the RareCapsNet architecture, *J* = 32 for all data sets. The number of cell types *T* (and hence *I* = *T*) can vary with *T* taking values between 11 and 15. Equally, the number of genes *K* supported by the particular data set can vary. In the experiments in which the algorithm here is used, we make use of data set 1 (mRNA + protein expression; CITE-seq^22^). According to this data set, *T* = *I* = 13 and *K* = 2000, see also the detailed specifications of the data sets in the Supplement.

In the following we outline an algorithm that, when provided with the trained MarkerCapsule network architecture and the training samples *S*, will output gene specific coupling coefficients 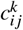. These 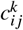 indicate how strongly gene *g*_*k*_ contributes to activating the link from primary capsule *j* to cell type capsule *i*.

The algorithm proceeds according to the following step:

1. For each training sample *s* ∈ *S* and each gene *g*_*k*_, *k* = 1, …, *K*, consider *s*^*k*^ defined by

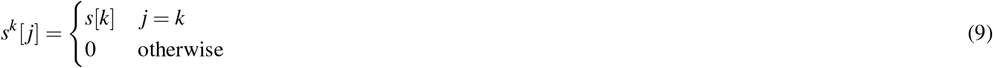

That is, *s*^*k*^ keeps *s*[*k*] from the original input of single cell *s* and has all other entries masked, that is, set to zero.

1. For each combination (*s, k*) of single cell *s* and gene *k*, we run *s*^*k*^ through the network. If additional data is considered, *s*^*k*^ is integrated with the additional data the usual way, before provided as input to the network.
2. Running *s*^*k*^ through the network includes executing the dynamic routing procedure, hence yields coupling coefficients 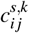 for all *i* = 1, …, *I, j* = 1, …, *J*
3. One averages the resulting 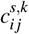 across samples assigning equal weight to each cell type *t*:

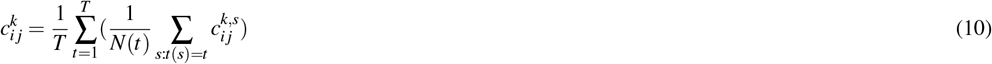
4. One outputs 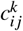 as the desired gene specific coupling coefficients.

### Algorithm for Computing Marker Genes through rare capsules

Given a particular cell type *i*, one determines the primary capsule *j* that yields the largest coupling coefficient *c*_*i j*_. Note that if several primary capsules *j* yield sufficiently large *c*_*i j*_, one can extend the analysis on all such *j*; for the sake of an analysis that gives rise to an unbiased comparison across cell types, we restrict ourselves to the primary capsule *j* that yields the larges *c*_*i j*_ for each *i*.

One then determines the gene specific coupling coupling coefficients 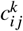 for all genes *g*_*k*_, *k* = 1, …, *K*, as per the algorithm described in Subsection . Subsequently, one determines the mean and the standard deviation of the 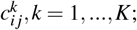, as can be seen in Figure 1 (for the combination of primary capsule 9 and cell type ‘CD16+Mono’, which exemplifies the situation for all selected combinations *j* and *i*), the empirical distribution of the 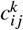 roughly follows a Gaussian distribution, with the exception of an heavy tail towards the right. The idea is to collect all genes that give rise to this heavy tail.

To do so in a sound way, we determine p-values 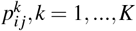 reflecting tail probabilities with respect to the Gaussian distribution the usual way. Subsequently, we sort the resulting 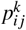 in ascending order (where the smallest 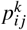 corresponds with the largest 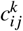) yielding

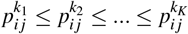

and determines

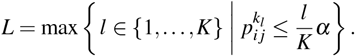

that is the largest *l* for which 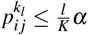, where *α* reflects the significance level at which one operates. Here, *α* = 0.05, which follows standard conventions. Overall, this procedure reflects the Benjamini-Hochberg procedure, which accounts for the necessary corrections due to multiple hypothesis testing^23–25^.

The set of marker genes *CM*_*i*_ for cell type *i* is then defined to be

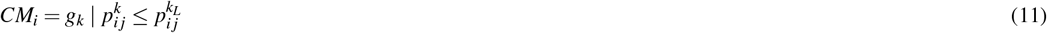

## Conclusions

In this work, we presented **RareCapsNet**, an interpretable capsule network–based framework for robust identification of rare cell populations from large-scale single-cell RNA sequencing data. By leveraging the dynamic routing mechanism of capsule networks, RareCapsNet explicitly models part–whole relationships between genes, primary capsules, and cell-type capsules, enabling reliable detection of rare cellular identities even under extreme class imbalance and high dropout settings in which conventional clustering-based approaches often fail.

A key strength of RareCapsNet lies in its interpretability. Through gene-specific coupling coefficients and statistically principled false discovery rate control, the framework not only detects rare cells but also identifies biologically meaningful marker genes associated with distinct capsule activations. Theoretical analysis establishes conditions for identifiability of rare cell types, robustness to class imbalance through routing normalization, and convergence of the routing procedure, thereby providing a sound mathematical basis for the observed empirical performance.

Extensive evaluation on simulated data and diverse real-world scRNA-seq datasets including Jurkat–293T mixtures, CBMC, PBMC68k, mouse brain, and cross-batch transfer settings demonstrates that RareCapsNet consistently outperforms state-of-the-art rare-cell detection methods in terms of accuracy, F1-score, and stability. Importantly, RareCapsNet maintains strong performance in poorly covered populations and enables cross-dataset generalization without retraining, highlighting its suitability for large, heterogeneous, and multi-batch single-cell studies.

While the current implementation focuses on supervised learning scenarios, RareCapsNet provides a flexible foundation for future extensions. Promising directions include semi-supervised and weakly supervised formulations, computational optimizations for large gene panels, and adaptation to multimodal and spatial transcriptomics data. Overall, RareCapsNet offers a scalable, interpretable, and theoretically grounded solution for rare cell discovery, advancing both methodological rigor and biological insight in single-cell transcriptomic analysis.

